# A Species-Specific Nematocide that Results in Terminal Embryogenesis

**DOI:** 10.1101/118141

**Authors:** Tess Renahan, Ray L. Hong

**Affiliations:** California State University, Northridge Department of Biology, 18111 Nordhoff Street Northridge CA, 91330-8303 U.S.A

**Keywords:** nematode, Pristionchus pacificus, insect pheromones, embryocide, eggshell, necromeny, host cues, pheromone, embryogenesis

## Abstract

Nematode-insect interactions are ubiquitous, complex, and constantly changing as the host and nematode coevolve. The entomophilic nematode *Pristionchus pacificus* is found on a myriad of beetle species worldwide, though the molecular dynamics of this relationship are largely unknown. To better understand how host cues affect *P. pacificus* embryogenesis, we characterized the threshold of sensitivity to the pheromone (Z)-7-tetradecen-2-one (ZTDO) by determining the minimum exposure duration and developmental window that results in *P. pacificus* embryonic lethality. We found early-stage embryos exposed to volatile ZTDO for as few as four hours all display terminal embryogenesis, characterized by punctuated development up to 48 hours later, with abnormal morphology and limited lumen formation. To determine if the pheromone arrests pre-hatching development by suffocating or permeabilizing the eggshells, we raised embryos under anoxic condition as well as examined eggshell permeability using the lipophilic dye FM4-64. We found that asphyxiating the embryos arrested embryogenesis in a reversible manner but did not phenocopy the effects of ZTDO exposure, whereas the ZTDO-induced disruption of embryogenesis did correlate with increased eggshell permeability. The effects of ZTDO are also highly specific, as other lipid insect compounds do not produce any detectable embryocidal effect. The high specificity and unusual teratogenic effect of ZTDO may be important in mediating the host-nematode relationship by regulating *P. pacificus* development.

**Summary Statement:** Insect-associated nematodes coordinate their development using host cues to “walk the line” between mutualism and pathogenesis. A host pheromone causes terminal embryogenesis by permeablizing the nematode eggshell.

## Introduction

Many parasites regulate their development to coordinate with that of their host species. In particular, the life cycle of parasitic nematodes includes one or more infective stages that respond to host cues, such as the dauer and pre-hatch juveniles. In most plant parasitic nematodes, the egg protects the pre-hatch juveniles but can still respond to the presence of hosts. For example, the hatching rate in plant parasitic nematodes *Heterodera spp* and *Globodera* spp. is significantly enhanced in the presence of root extracts (Byrne et al., 2001; Ibrahim and Haydock, 1999; Tefft and Bone, 1985). However, the coordination between the development of insect parasitic nematodes and their insect hosts is not well documented. In particular, the entomophilic nematodes that remain non-pathogenic on specific hosts may reveal novel signaling mechanisms necessary to regulate developmental progress between nematodes and their insect hosts.

In a type of association known as necromeny, the nematode *Pristionchus pacificus* and related nematodes live on various species of beetles and wait until their hosts die before resuming reproductive development to feed on the microorganisms growing on the host carcasses (Herrmann et al., 2006a; Herrmann et al., 2006b; Manegold and Kiontke, 2001; Sudhaus, 2009). However, since entomphilic nematodes can continue to evolve new relationships with potential pathogenic bacteria to infect insect hosts (Dillman et al., 2012; Ye et al., 2010; Zhang et al., 2008), it is not known how *Pristionchus* and related nematodes maintain their strictly necromenic and non-pathogenic interaction with their hosts. It is possible that the more species-specific relationship found in certain *P. pacificus* populations have mechanisms that prevent reproductive development on living hosts, such as the association between *P. pacificus* and the oriental beetle *Exomala orientalis* (Herrmann et al., 2007). Such mechanism may depend on specific host cues that can regulate a durable but nonpathogenic necromenic relationship. In particular, the sex pheromone of the oriental beetle, (Z)-7-tetradecen-2-ol (ZTDO) (Zhang et al., 1994), is a chemical attractant for adult nematodes as well as a nematocide that paralyzes early larval stages, prevents dauer larvae exit, and inhibits egg hatching (Cinkornpumin et al., 2014). Most surprisingly, unlike other natural compounds so far described, ZTDO is a volatile nematocide that acts without direct contact. Because responses to ZTDO depend on the developmental stage in the *P. pacificus* life cycle, ZTDO represents a key semiochemical important for the coordination of the necromenic relationship between the nematodes and the beetles (Chaisson and Hallem, 2012; Leal, 2005; Okumura et al., 2013; Ruther et al., 2002).

Given the well-known ability of the nematode eggshell to resist chemical and biological assaults, the ability of ZTDO to elicit novel embryonic lethal phenotypes is especially intriguing. The nematode eggshell is one of the most resistant biological structures and a major barrier against chemical and enzymatic nematocides. The nematode eggshell may consist of up to five layers, but the stereotypic nematode eggshell is a trilaminar structure comprised of an inner lipid layer, a middle chitinous layer, and an outer vitelline layer (Rappleye et al., 1999; Wharton, 1980). These layers assemble soon after fertilization of the oocyte by the sperm to prevent polyspermy as well as to protect the developing embryo from the environment. Nevertheless, the eggshell is still permeable to oxygen as well as water. Environmental factors can also penetrate the eggshell, sometimes in a species-specific manner. For example, the nematophagous fungi *Paecilomyces lilacinus* produces chitinases that degrades the outer two layers of the eggshell of the plant parasitic nematode *Meloidogyne javanica,* as well as proteases that destroy the inner lipid layer (Khan et al., 2004). Other filamentous fungi also use chitinases to attack the eggs of the parasitic nematode *Ascaris lumbricoides* during its free-living stage in the soil (Kunert, 1992). Even plant defense compounds, such as trans-2-hexenal, can significantly reduce the hatching rate of the pinewood nematode *Bursaphelenchus xylophilus* after a 12- hour treatment (Cheng et al., 2016). Thus despite a remarkably resistant structure, the nematode eggshell represents both a regulator for host-parasite communication as well as a valid target for host derived anthelmintics.

While the chitinous layer provides rigidity, the lipid layer is widely perceived to act as a permeablility barrier against most chemicals. Recent studies in the compost-dwelling *Caenorhabditis elegans* revealed a fourth layer between the embryo and the trilaminar eggshell that confers impermeability but is distinct from the lipid layer (Olson et al., 2012). Genetic analysis suggests that this permeability barrier consists of ascarosides with lipid side chains. Although the eponymous ascarosides are found in the lipid layer of eggshells of Ascarids, the conservation of this inner permeablility barrier in Ascarid species and other nematodes has not yet been examined. The permeability barrier in *C elegans* can be compromised by a small compound, C22, that results in weakened eggshells sensitive to minor osmotic changes and light pressure. C22 however, acts specifically on the oocyte-to-embryo transition in *C. elegans* but not in other *Caenorhabditis* species, suggesting that the permeability barrier evolves quickly among nematodes.

In this study, we show that a minute amount of the sex pheromone ZTDO from the oriental beetle host is a potent embryocide that can permeablize and perhaps also penetrate the *P. pacificus* eggshell. Exposure of early 2-8 cell embryos to the volatile form of ZTDO elicits a potent lethal effect on embryos that derails normal development and permanently prevents embryos from hatching. More curiously, most embryos continue to develop long after treatment has ended, with around half of the embryos developing muscles, gut, and pharynx-like structures well past the normal hatching time of 24 hours. This embryonic lethality varied among global *P. pacificus* strains, with the strongest effect found in a Japanese strain isolated from the oriental beetle, and is correlated with ZTDO-induced eggshell permeability. However, eggshell permeability alone does not account for all of the ZTDO-induced lethality, which may suggest that zygotic factors also mediate ZTDO susceptibility.

## Materials and Methods

### Nematode Strains

All *P. pacificus* and *C. elegans* strains were maintained on 6 cm NGM plates seeded with OP50 and kept at 20°C as (Hong et al., 2008). The following *P. pacificus* strains were used: PS312 (California wild-type, synonymous with RS2333), PS1843 (Port Angeles, WA), *Ppa-obi-1(tu404)I, csuls01[Ppa-obi-1p::gfp; Ppa-egl-20p::rfp; PS312* gDNA]X, RSB001 (La Reunion), RS5278 (Bolivia), RS5419 (La Reunion), RS5186 (Japan), RS5194 (Japan), and RLH163 (Mar Vista, CA). Wild-type N2 *C. elegans* was used. *P. pacificus* populations include both hermaphrodites and ~1% males and have a life cycle of four days.

### Lid-drop assay

The ZTDO assay was used to determine both the minimum lethal exposure duration of wild-type PS312 embryos and ZTDO susceptibility of other strains. To synchronize embryos, approximately 20 well-fed, young gravid hermaphrodites were isolated on 6 cm plates (T3308; Tritech, Los Angeles, CA)with 100 μl of OP50 and then removed after an hour. For ZTDO treatment, 10 μl of 0.5% (Z)-7-tetradecen-2-ol (ZTDO; Bedoukian Research, Danbury, CT) diluted in 100% ethanol (v/v) was added to the underside of the plate lids and sealed with Parafilm^®^. Given the dimensions of the cylindrical plate (r= 2.5 cm, h=0.5 cm), the volume encompassing the embryos above the agar is approximately 9.82 cm^3^. Thus the addition of 0.05 μl ZTDO results in at least 5.1 x 10^6^ dilution, not accounting for any ZTDO uptake by the agar. (E)-11-tetracenyl acetate (EDTA; Bedoukian Research, Danbury, CT) and Methyl tetradecanoate (myristate; Sigma-Aldrich, St. Louis, MO) were diluted to 0.5% in ethanol and administered the same way as ZTDO. To stop the treatment, Parafilm and lids were removed after exposure and replaced with new lids. To determine hatching rates, the number of eggs was counted after synchronization (time 0) and unhatched embryos were counted at screening time (26 hours or 48 hours). N(n)=number of experimental trials performed with control on different days (total sample size of nematodes). Trials with unusually high arrest rates in control embryos were omitted.

### Microscopy

For determining the hatching rate, eggs and larvae were score using the Leica S6E dissecting microscope (Wetzlar, Germany). For FM4-64 staining, embryos were viewed using Differential Interference Contrast (DIC) on a Leica DM6000 upright microscope. Embryos were fixed in a 50:49 (egg salts:dH_2_0) solution on slides inside a small ring of Vaseline^®^ and covered with 22 x 22 mm coverslips on 3” x 1” x 1.0 mm slides. To determine permeability, embryos were fixed in a solution of 1:100 (FM4-64:egg buffer) from a 300 μM FM4-64 stock solution (Molecular Probes, T3166, Eugene, OR). Images were captured using the exposure time of one second and at an intensity level of two on the Leica Application Suite (LAS). Images were cropped using Adobe Photoshop CS6 (San Jose, CA).

### Anaerobic chamber

For the anaerobic experiments, a sealed jar and Oxoid^™^ AnaeroGen^™^ sachets (Thermo Fisher Scientific, AN0035A, Waltham, MA) were used. Oxygen levels were confirmed using Oxoid Anaerobic Indicator strips (Thermo Fisher Scientific, BR0055). An hour was added to anaerobic exposure times to allow oxygen levels to reach below 1%. Carbon dioxide levels reach between 9-13%, and no nitrogen was produced as a side-product from the sachets.

### Statistical analysis

Statistical tests were analyzed using GraphPad Prism (La Jolla, CA) and Microsoft Excel.

## Results

### ZTDO is a potent embryocide

Our previous study showed that *P. pacificus* early-stage embryos exposed for two days to minute amounts of the oriental Beetle pheromone (z)-7-tetradecen-2- ol (ZTDO) severely retards embryonic development and almost all embryos remain at a pre-tadpole stage two days later, whereas most embryos exposed to the ethanol vehicle hatch approximately one day later (Cinkornpumin et al., 2014). To delineate the earliest embryonic stage most sensitive to ZTDO, we focused on its embryocidal effect on the first half of embryonic development using defined exposure durations, rather than continuous exposure. Because exposing embryos to less than 0.5% ZTDO for two days results in some embryos hatching, rather than varying the concentration of ZTDO we sought to determine the lethal time and susceptibility window necessary for 10 μl of volatile 0.5% ZTDO to prevent eggs from hatching after 24 hours. In the lid-drop assay, staged embryos are exposed to a volatile source of 0.5% ZTDO under the lid of a culturing plate for a defined time, after which the lid is replaced with a new one to terminate the treatment (Fig. 1). This is approximately a 5.1 x 10^6^ fold ZTDO dilution in the headspace above the assay arena. Unlike *C. elegans* that tend to hold onto eggs in its uterus, wild-type *P. pacificus* PS312 young adult hermaphrodites typically hold on to only 2 eggs in the uterus. Eggs are usually laid at the 2-cell stage, though embryos can be expelled from the uterus as old as the 8-cell stage. Embryos synchronized to an hour range from 2 to 16 cells at 20°C (Fig. 2A-E) and *P. pacificus* embryogenesis requires 24 hours at 20°C, compared to 18 hours for *C. elegans* (Felix et al., 1999; Vangestel et al., 2008). The proportion of unhatched eggs, including unfertilized oocytes and spontaneously arrested embryos, are counted after 26 hours *ex utero.* The additional two hours is to accommodate the time needed to view the embryos. Roughly 95% of ethanol-treated control eggs hatch after 26 hours (Fig. 2). ZTDO exposure on gravid hermaphrodites has a detectable but statistically insignificant effect on lowering the egg laying rate, but has no effects on the hatching rate of embryos exposed *in utero* (data not shown).

**Fig. 1.**
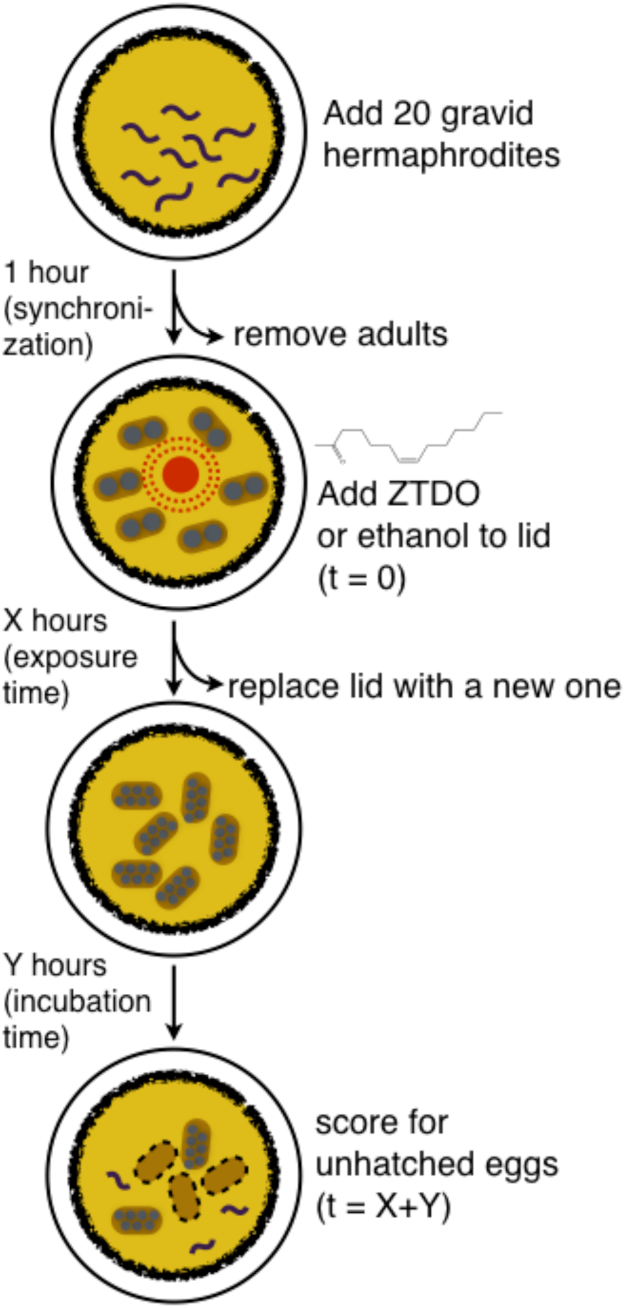
The lid-drop assay for ZTDO sensitivity. Synchronized 1-hour *ex utero-* old embryos were exposed to 10 μl of either 0.5% ZTDO (z-7-tetradecen-2-ol) or control (ethanol) suspended from plate lids.

**Fig. 2.**
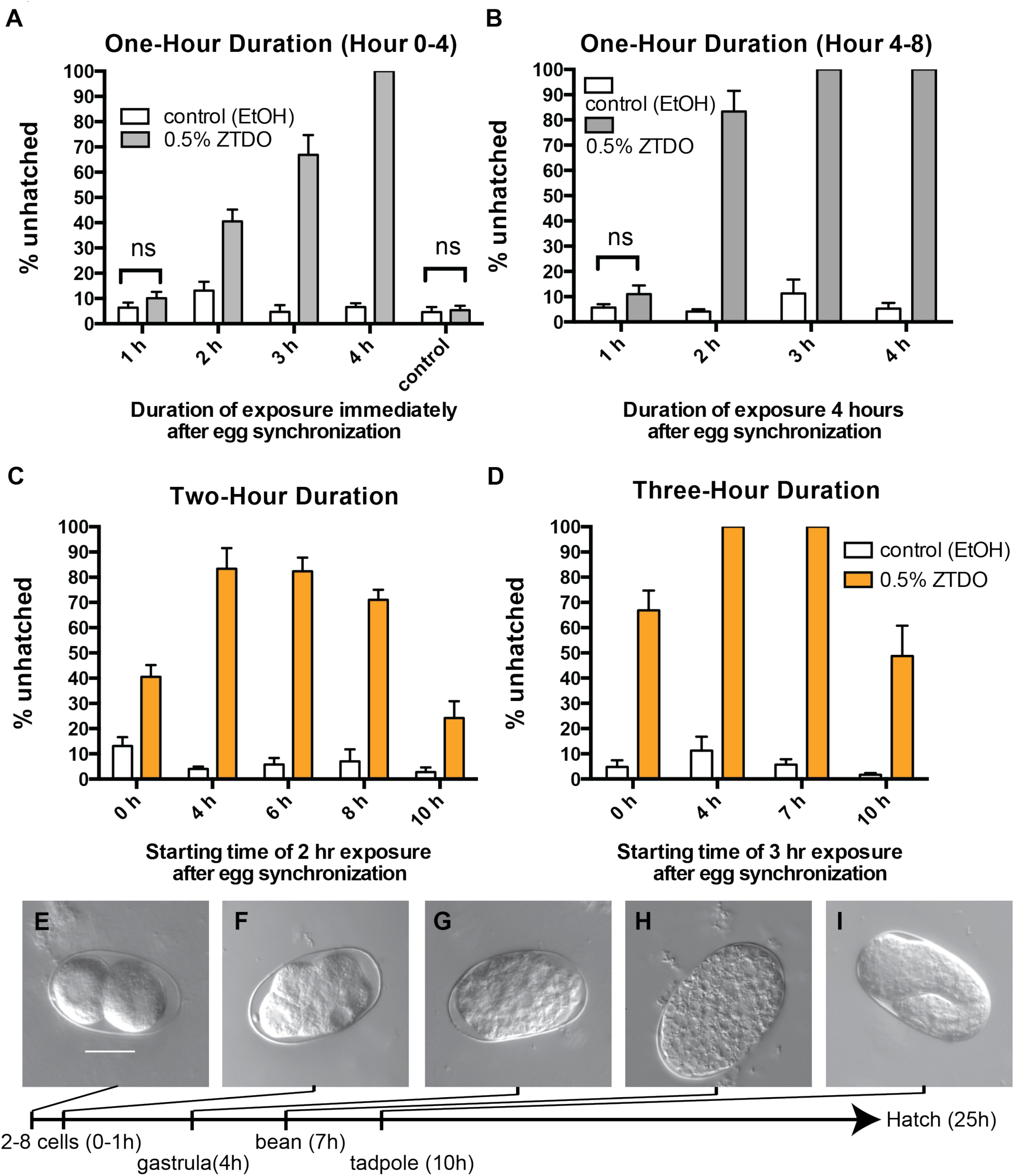
ZTDO sensitivity *ex utero* depends on exposure duration and embryonic stage. (A–E) Representative embryonic stages of untreated *P. pacificus* PS312 eggs *ex utero* at 20°C. Eggs are laid as early as the 2-cell stage and as late as the 8-cell stage. After an hour of synchronization, just prior to ZTDO exposure, the oldest embryo is 16-cell. Most embryos arrest during gastrulation and the bean stage. 10 μl of 0.5% ZTDO or ethanol was suspended from each plate lid. The scale bar of 20 μm in (A) represents the same magnification for all images.(F) Early embryos up to four hours *ex utero* show developmental arrest with increasing exposure time to ZTDO. To determine the possible effects of residual ZTDO, untreated embryos were added to NGM plates previously exposed to ZTDO for four hours (control). N(*n*) from left to right: 5(147), 5(132), 10(420), 10(432), 10(432), 10(406), 10(476), 10(454), 10(486), 10(416). 5) Older embryos between 4-8 hours *ex utero* old show higher sensitivity to ZTDO. N(*n*) from left to right: 5(296), 5(309), 5(305), 5(383), 5(262), 5(308), 5(355), 5(287). (H) When exposed to ZTDO for only two hours, 4-6 hour old embryos show higher rates of developmental arrest impared to just laid and 8-10 hour old embryos. 2-hour exposure data is the same as in 2F and 2G started at 0 and 4^th^ hour *ex utero.* N(n) from left to right: 10(432), 10(406), 5(305), 383), 5(291), 5(336), 5(270), 5(317), 5(239), 5(265). (I) When exposed to ZTDO for three hours, 4-7 hour old embryos showed even higher rates of developmental arrest compared to st laid and 10 hour old embryos. 3-hour exposure data is the same as in 2H and 2I started at 0 and 4^th^ hour ex *utero.* N(*n*): 10(476), 10(454), 5(262), 5(308), 5(310), 5(298), 5(283), 5(373). error bars denote standard error of the mean.

We found that the earliest response to ZTDO occurs when synchronized embryos are exposed for at least 2 hours, in which 40% of the embryos arrest. These embryos are between 120-180 minutes old and correspond to the start of gastrulation at the 28-cell stage, approximately 185 minutes after fertilization (Vangestel et al., 2008). Lethality reaches 100% when exposure is extended to 4 hours (Fig. 2F). As a control, untreated embryos added to NGM OP50 plates previously exposed to ZTDO for four hours do not show any responses to residual ZTDO. Thus, the removal of the lid is sufficient to terminate the volatile ZTDO treatment. The ZTDO embryocidal effect is even stronger in older embryos that are exposed to the same dose of ZTDO between 4-8 hours after synchronization, in which a 2-hour exposure resulted in 80% embryonic lethality, and a 3-hour exposure resulted in total lethality (Fig. 2G). These results show that the lethal time for a 5.1 million dilution of ZTDO is just two hours rather than only after two days of exposure (Cinkornpumin et al., 2014), and that the later 4-8 hour period *ex utero* is more sensitive than the earlier 0-4 hour period.

To determine if a shorter exposure time to achieve 100% lethal arrest is possible in other 2-3 hour windows within the first 13 hours of development *ex utero,* we exposed embryos to ZTDO for two hours in two-hour time windows and found that lethality peaks at 80% (Fig. 2H). However, extending ZTDO exposure to three hours between 4-7 hours and 7-10 hours *ex utero* results in the terminal arrest of all embryos, suggesting that the period between the gastrula and tadpole stage is much more sensitive to volatile ZTDO than the periods bracketing it (Fig. 2I). In *C. elegans,* the period leading to the tadpole stage is characterized by cell proliferation (Sulston et al., 1983), therefore it is likely that in *P. pacificus* ZTDO may interfere directly or indirectly with the process of cell division. In very rare circumstances, usually in association with bacterial contamination that results in thick lawns, ZTDO-exposed embryos show delayed hatching 48 hours *ex utero*. Because the earliest embryos that were exposed to ZTDO for four hours all arrest, we performed subsequent characterizations using this standard regiment: 1-2 hours of embryo synchronization, followed by 4 hours of ZTDO exposure, and scored for hatching success after 26 hours *ex utero.*

### Natural variation in ZTDO sensitivity

Given that *P. pacificus* populations are found to be associated with several different beetle species globally (Herrmann et al., 2006a; Herrmann et al., 2006b; Rodelsperger et al., 2014) and that chemotaxis attraction towards ZTDO differ in various *P. pacificus* strains (Hong et al., 2008; Koneru et al., 2016), we surmised that the embryocidal sensitivity to ZTDO may also vary across different strains. Using the lid drop assay with synchronized embryos exposed to 0.5% ZTDO for four hours, we observed that strong differences in embryocidal sensitivity correlated with geographic origins. Like the reference strain PS312 from California, the strains from the Pacific Rim (RLH163 California; PS1843 Washington; RS5194 and RS5186 Japan) have highly ZTDO-sensitive embryos (Fig. 3). In contrast, a strain from Bolivia (RS5278) and three strains from La Reunion Island in the Indian Ocean (RS5419, RS5413, RSB001) all show 40% or higher of embryos hatching in the presence of ZTDO. *C. elegans* N2 is insensitive to ZTDO at the concentration tested, even after two days of exposure (Cinkornpumin et al., 2014). Interestingly, host origin is only weakly correlated with ZTDO sensitivity. The sensitive strain RLH163 was isolated from the masked chafer *Cyclocephala pasadenae,* while the RS5194 and RS5196 Japanese strains were isolated from the Oriental beetle *Exomala orientalis.* The four resistant strains from Bolivia and La Reunion were isolated from four different species of beetles (Rodelsperger et al., 2014). Since the oriental beetle pheromone is the only *P. pacificus* host compound commercially available, it is difficult to test if the correlation between ZTDO sensitivity and geography, but not host species, is due to overlap in pheromone structure between *Cyclocephala* and *Exomala* in sensitive strains, or rapid host switching in resistant strains (McGaughran et al., 2013).

**Figure 3.**
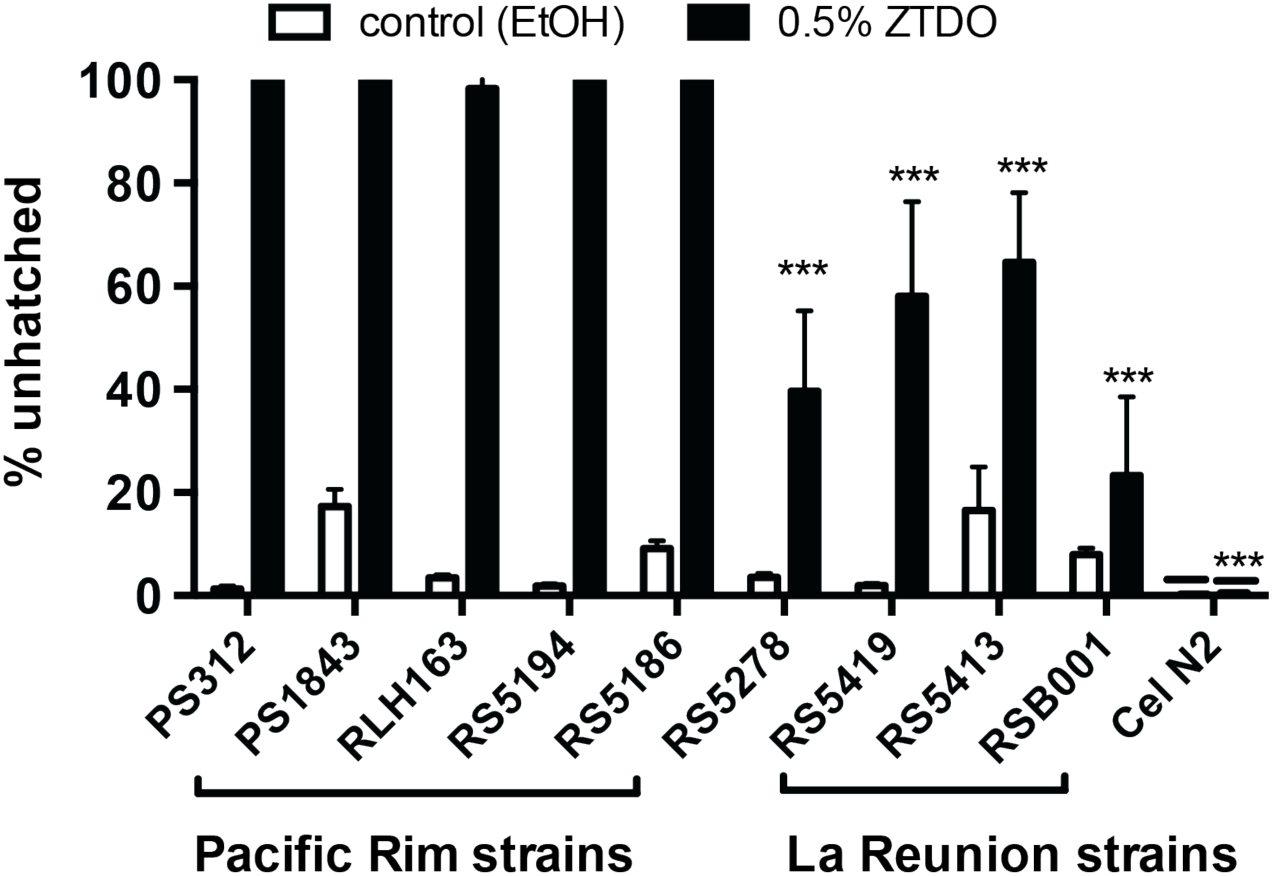
Natural variation in *P. pacificus* responses to the embryocide ZTDO. Early-stage embryos were exposed to 10 μl of 0.5% ZTDO suspended from plate lids for four hours. Strains from the Pacific Rim - PS1843 (Washington), RLH163 (California), RS5194 (Japan), RS5186 (Japan) - resemble the reference strain PS312 (California). Strains from La Reunion (RS5419, RS5413, RSB001) and Bolivia (RS5278) are significantly less sensitive to ZTDO. *C. elegans* N2 do not respond to 0.5% ZTDO. All differences between ethanol control and 0.5% ZTDO were significant except for RSB001 and N2, as determined by paired t-test (P<0.05). Significant difference to PS312 was determined by one-way ANOVA followed by Dunnett’s (***P<0.0001). Error bars denote standard error of the mean. N(n): 10(926), 10(948), 15(2,894), 15(3,771), 15(1,899), 15(2,246), 15(1,678), 15(1,514), 15(1,482), 15(2,194), 15(1,965), 15(2,640), 15(2,735), 15(2,844), 15(2077), 15(2011), 15(3,497), 15(3,275), 15(4,335), 10(3,098).

### Anoxia does not phenocopy the effect of ZTDO

The development of the nematode embryo within the egg is aerobic and requires an external source of oxygen passing through the eggshell. Given the much larger molecular structure of ZTDO compared to those of water and oxygen, and the presumption that nematode eggshells are impervious to most small molecules (Carvalho et al., 2011; Chitwood and Chitwood, 1974), we wondered if rather than penetrating the eggshell, ZTDO acts to arrest development by coating the eggshell and blocking essential gas exchange. To see if oxygen deprivation would phenocopy the embryocidal effect of ZTDO, we incubated synchronized *P. pacificus* embryos in an anaerobic chamber for five hours and then scored for hatching embryos at 24 hours *ex utero. C. elegans* embryos were also deprived of oxygen for five hours before resuming normal incubation on the plate and scored at 18 hours *ex utero* (*C. elegans* has a shorter embryogenesis). However, the deprivation of oxygen for five hours neither replicated the ZTDO embryo phenotype nor delayed hatching (Fig. 4A). Moreover, when we exposed synchronized *P. pacificus* embryos to ethanol or left them alone for three hours, and then incubated them under anoxic condition for 19 hours, most embryos hatched 24 hours later, *i.e.* around 50 hours *ex utero.* In contrast, no embryos hatched after 50 hours when they were exposed to ZTDO prior to anaerobic treatment (Fig. 4B). Almost none of the ZTDO-exposed embryos hatched even three days after normal hatching time. These results suggest that like *C. elegans* embryos, *P. pacificus* embryos also undergo anoxia-induced arrest that delay development and hatching (Padilla and Ladage, 2014). However, unlike anoxia-induced arrest, the ZTDO-induced arrest is almost always irreversible in *P. pacificus* embryos at the minimum exposure regiment necessary to prevent hatching.

**Fig. 4.**
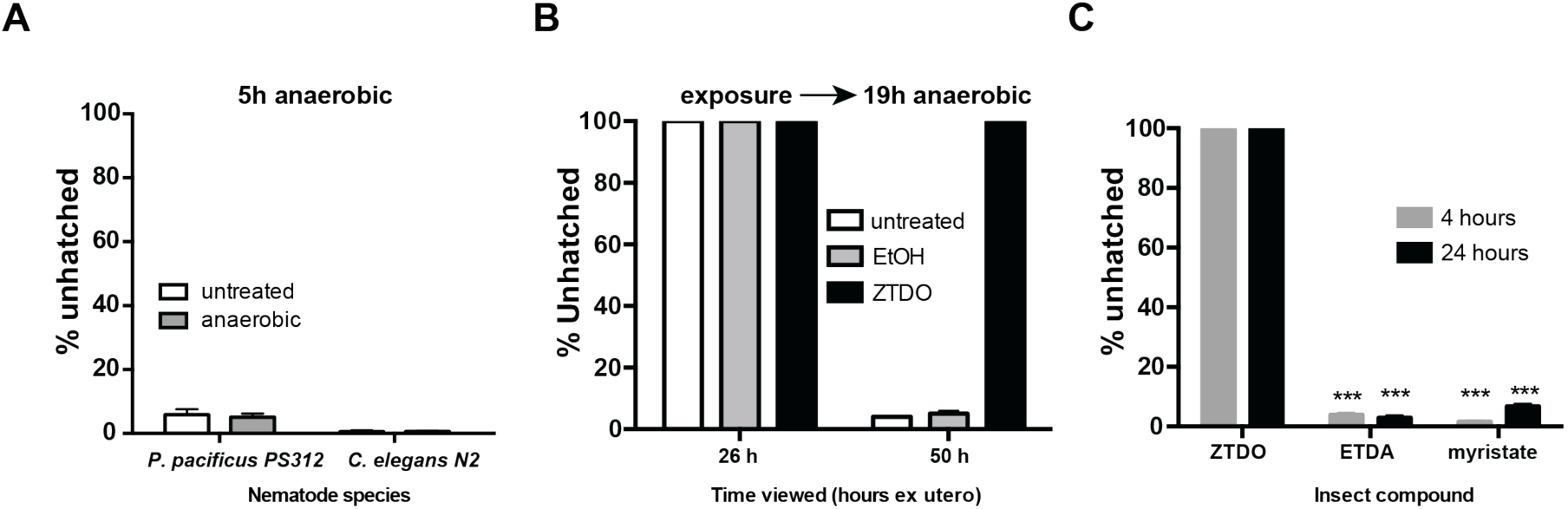
Anaerobic condition and other insect compounds do not phenocopy the effects of ZTDO. (A) Synchronized *P. pacificus* and *C. elegans* embryos were deprived of oxygen at 20°C for five hours, then allowed to recover with oxygen for 24 or 18 hours, respectively. N(n)=3(512), 3(807), 3(707), 3(651). (B) Synchronized *P. pacificus* embryos were left alone (untreated) or exposed to ethanol or ZTDO for 3 hours, followed by 19 hours without oxygen. The anaerobic condition delayed egg hatching but ZTDO pre-treatment arrested embryogenesis permanently. N(n)=4(611), 5(654), 5(650), (C)P. *pacificus* PS312 was exposed to 10 μl of 0.5% of the insect compound suspended from plate lids for four and 24 hours. E-(11)-tetradecenyl acetate (ETDA). Significant differences in response to the compounds were determined by two-way ANOVA (***P<0.0001). N(n)=15(1422), 15(1690), 15(1177), 15(1090), 15(2218), 15(1496). Error bars denote standard error of the mean.

Finally, to investigate whether other insect-derived, long-chain hydrocarbon compounds attractive to *P. pacificus* can also act as embryocides, we performed the lid-drop assay using E-(11)-tetradecenyl acetate (ETDA), a moth sex pheromone from *Spodoptera littoralis,* and methyl myristate, an allomone from *Musca* flies (Hong and Sommer, 2006). We found that neither compound prevents *P. pacificus* embryos from hatching into healthy J2 larvae (Fig. 4C). Therefore, ZTDO is a species-specific embryocide.

### Aberrant and incomplete embryonic development

Approximately a quarter (26%) of ZTDO-exposed early embryos stop development at the bean stage, while more than half arrest in later stages that exhibit partial lumen formation and muscle contraction. The “peanut” stage, containing two unequal sized lobes of cells, is a unique stage for ZTDO-treated embryos that seems to have initiated elongation but does not reach the tadpole stage (Fig. 5A). Interestingly, although muscle contraction does not start until after the tadpole stage in *C. elegans,* some ZTDO-treated peanut stage *P. pacificus* embryos exhibit movement. Sporadic twisting and whole-body bending movements can also be seen in the later 2-fold and 3-fold stages (Fig. S1A-C). More intriguingly, lumen-like cavities appear in the peanut, 2-fold, and 3-fold stages, with increasing proportion of these lumens occurring in these stages found at 48 hours *ex utero* (Fig. 5B-E). Specifically, the proportion of peanut-stage embryos with lumen quadrupled between 26-48 hours, while lumen formation doubled in 2-fold and 3-fold stage embryos during the same period. Lumens near the extremes of the aberrant embryos resemble partial pharynges (Fig. 5B, D), while lumens near the center of the embryos resemble partial intestines (Fig. 5C, E). One major difference in *P. pacificus* from *C. elegans* is the roughly 80-minute delay in the first E cell division after the first MS cell division, around 70 minutes after fertilization (Vangestel et al., 2008). This temporal separation between the MS and E divisions may help to explain the variation in ZTDO phenotypes— the MS cell divides shortly after starting the ZTDO treatment for some embryos, while the E division always occurs long after the start of the ZTDO treatment for all embryos. We surmised that in our assays, the stem cell of the endoderm lineage, the E cell, is exposed to ZTDO longer than the MS mesoderm precursor cell. The ability to contract muscles and coordinate cell movements to create cavities suggests at least partial developmental determination in these moribund embryos, though it is unclear if certain cell lineages are more susceptible than others due to cell identity or length of exposure.

**Fig. 5.**
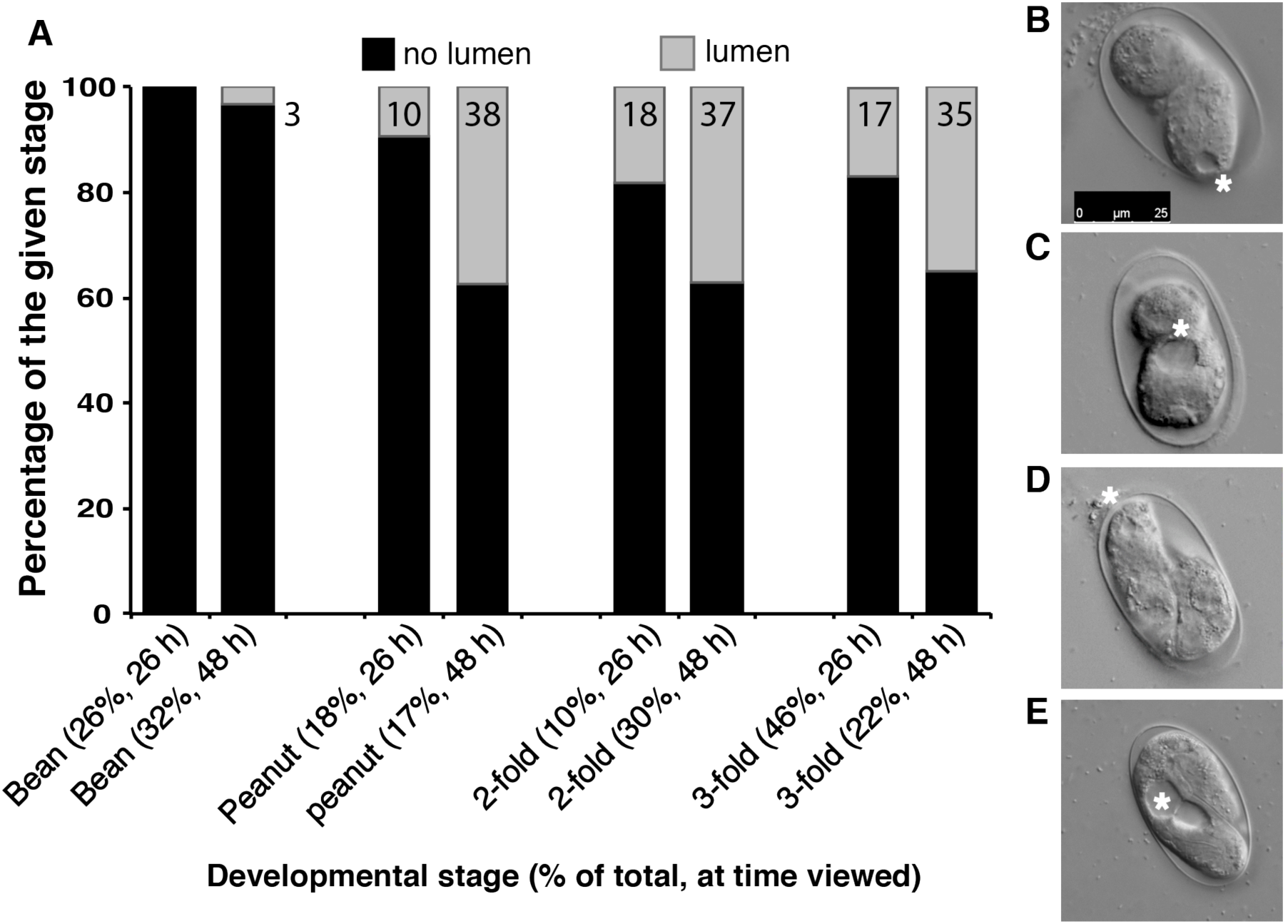
Aberrant development in ZTDO-exposed *P. pacificus* embryos. Representative images of early-stage embryos exposed to 10 μl of 0.5% ZTDO suspended from plate lids for four hours immediately following egg synchronization. The formation of a lumen-like structure for all embryonic stages increased with incubation time beyond the normal 26 hours. (A) The ‘bean’ stage is the least developed of the zombryo stages, representing 26% and 32% of the total viewed at 26 and 48 hours ex utero. (B) A ‘peanut’ stage embryo with an anterior lumen viewed at 48 hours *ex utero.* (C) A ‘peanut’ stage embryo with a mid-body lumen viewed at 48 hours *ex utero.* (D) A 2-fold stage embryo with an open anterior lumen viewed at 26 hours *ex utero.* (E) A 3fold stage embryo with a mid-body lumen viewed at 26 hours *ex utero.* *denotes lumen. The scale bar of 25 μm in (B) represents the same magnification for all images. N(n)= 3(115) (26 hours), 3(93) (48 hours).

Although we did not follow the developmental trajectories of individual embryos, but rather sampled the exposed population for the two time periods in three separate experiments, we speculate that these developmentally advanced “zombryos” attempt to continue cell differentiation and form the alimentary system as well as coordinated muscle contraction even when body elongation cannot be properly carried out and when embryogenesis is extended to twice the usual duration. Curiously, zombryos viewed at 48 hours *ex utero* show decreased proportion of 3-fold embryos compared those viewed at 26 hours. One explanation could be that the 3-fold embryos visible at 26 hours contracted to resemble 2-fold embryos by 48 hours. Unlike *C. elegans, P. pacificus* embryos transition to the pre-hatching J1 larvae inside the eggshell and molt just prior to hatching as J2 larvae (Lewis and Hong, 2014). However, the pre-hatching molt is unlikely to contribute to the hatching phenotype, since the embryos were exposed during early development before the gastrula stage. Interestingly, we found that some 2-fold stage zombryos exhibit *Ppa-obi-1p::gfp* cell expression characteristic of pattern found in the epidermis of untreated pre-hatching J1’s (Cinkornpumin and Hong, 2011)(Fig. 6C). We interpret the expression of seam cell markers to mean partial differentiation of the epidermal lineage, despite retarded overall development. Thus for the small proportion of the ZTDO-treated embryos that become zombryos, partial development continues but not enough to trigger hatching or promote survival outside the eggshell. Even the more advanced 48-hour old 3-fold stage zombryos do not survive on NGM media when we intervened to pop open the eggshell by compressing the zombryos between an agar pad and a cover slip. The variability and delay in the zombryo phenotype suggest more than one cell type may be targeted by ZTDO..Because the punctuated and incomplete embryogenesis does not seem reversible or recoverable, it is not clear when these zombryos actually seize living as embryos.

**Fig. 6.**
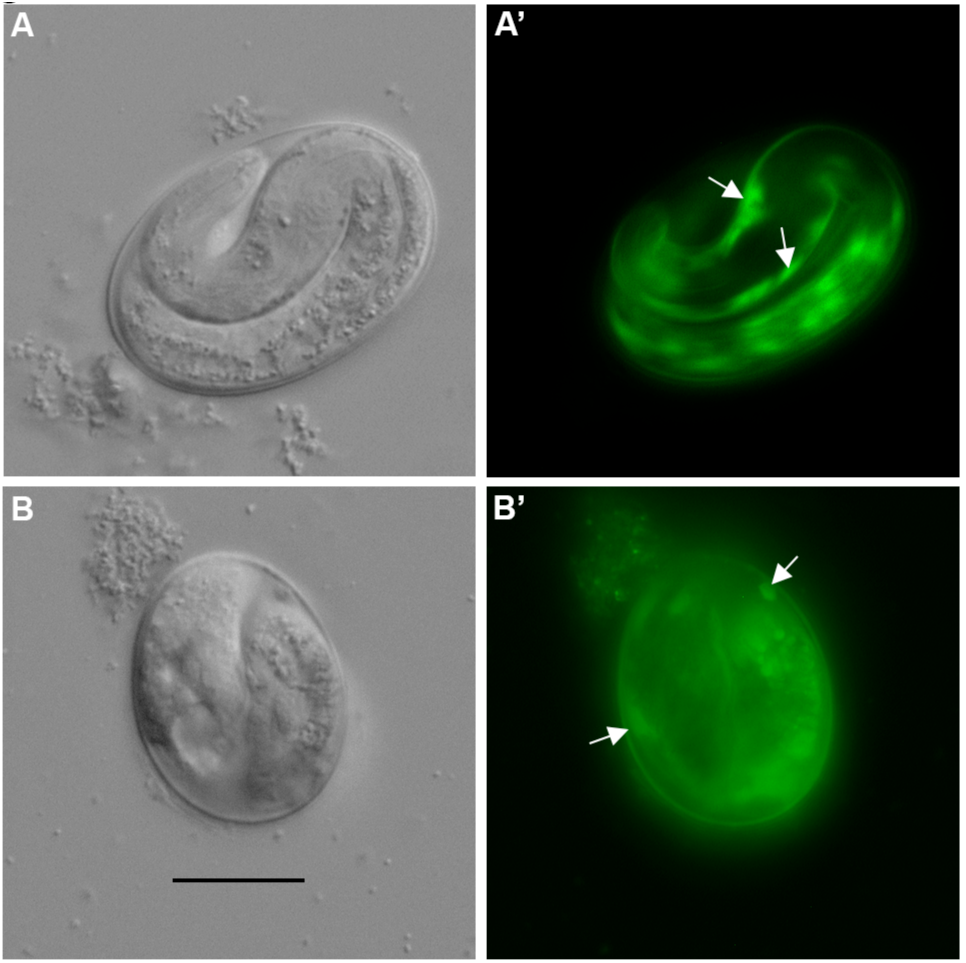
Expression of a J1 stage marker in arrested *P. pacificus* embryos. (A, A’) Seam cell expression *Ppa-obi-1p::gfp* (arrows) in untreated 24 hour-old embryos. (B, B’) Similar seam cell expression (arrows) can be observed in ZTDO-induced arrested 24 hour-old embryos. The scale bar of 20 μm in (B) represents the same magnification for all images.

### ZTDO increases eggshell permeability

Given that anoxia and other insect-derived pheromones do not produce embryonic arrests similar the ZTDO-exposure phenotype, we wondered if ZTDO enters through the eggshell to target embryonic tissues. We speculated that since we expose ZTDO to eggs after fertilization and *ex utero,* ZTDO is unlikely to interfere with eggshell assembly (Olson et al., 2012), but rather with eggshell function or integrity. Previous studies demonstrated that laser-permeabilized wild-type *C. elegans* embryos enable the fluorescent lipophilic dye FM4-64 to label the plasma membrane and endocytic structures (Rappleye et al., 1999). We found that FM4-64 strongly stains embryonic membranes of early *P. pacificus* embryos exposed to ZTDO for four hours, but only weakly stains the eggshell in the ethanol-exposed control (Fig. 7A-B). This ZTDO-induced eggshell permeability appears to be irreversible, as FM4-64 also stains later 2-fold stage zombryos 20 hours after ZTDO exposure ended, in contrast to the lack of staining in the control J1 pre-hatch larvae (Fig. 7C-D). ZTDO-treated embryos result in expanded peri-embryonic space, possibly due to shrinking of embryonic tissue. ZTDO-exposed embryos also shrink relative to the eggshell on the culturing NGM medium without mounting on agar, and exposure of embryos immersed in *C. elegans* Blastomere Culture Media (Shelton’s Media) did not ameliorate the ZTDO effect (data not shown)(Skop et al., 2001), suggesting that a more permeable eggshell contributes only in part to the increased osmotic sensitivity.

**Fig. 7.**
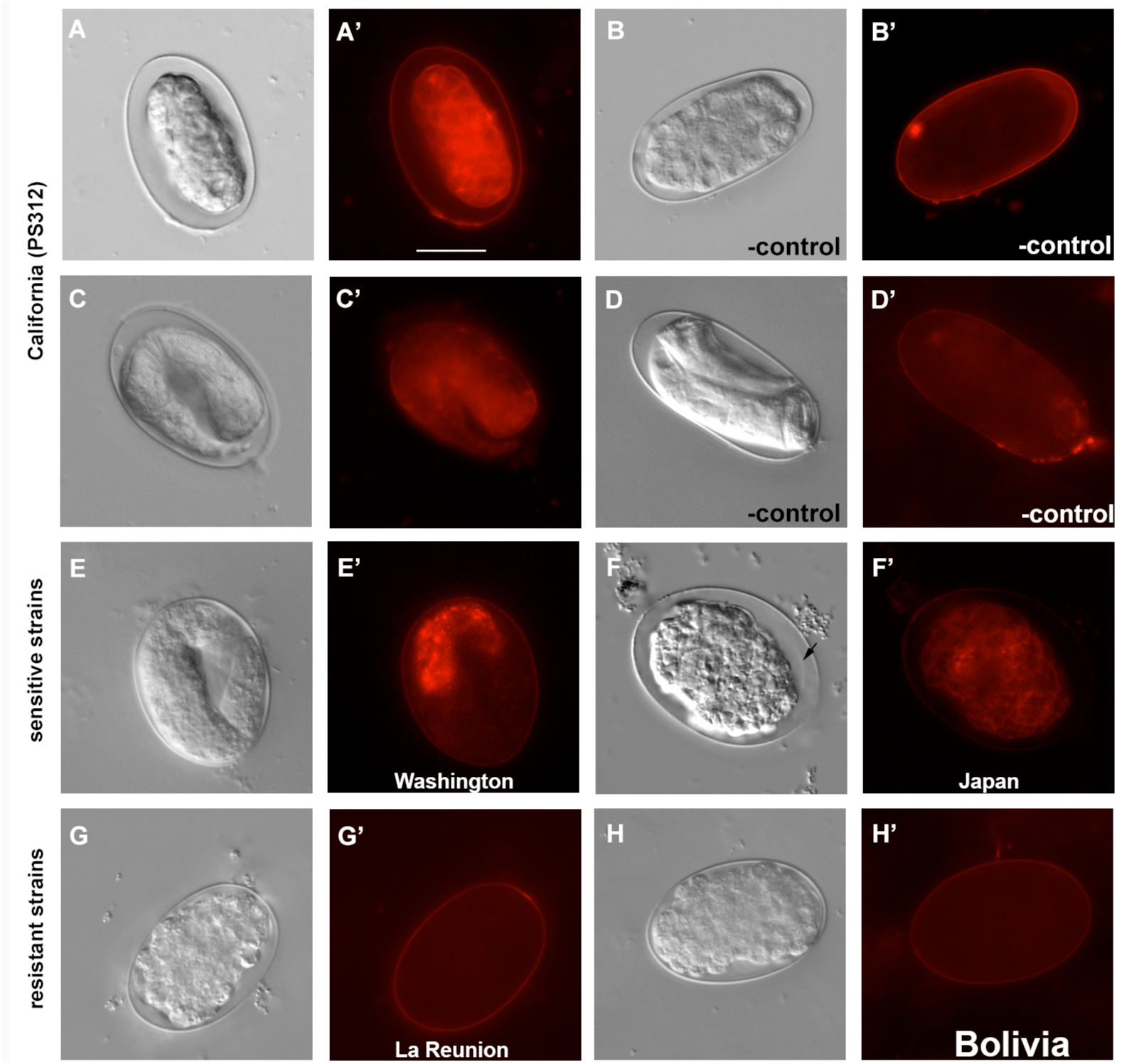
FM4-64 membrane staining of *P. pacificus* embryos exposed to volatile ZTDO. Representative images of early embryos exposed to 10 μl of 0.5% ZTDO suspended from plate lids for four hours. (A, A’) Wildtype PS312 exposed to ZTDO viewed immediately after treatment show stained embryonic tissue due to a permeable eggshell. (B) Wildtype PS312 exposed to ethanol control viewed immediately after treatment show only weak eggshell staining due to an impermeable eggshell. (C, C’) Wildtype PS312 exposed to ZTDO viewed ~20 hours after treatment. (D, D’) Wildtype PS312 exposed to ethanol vehicle control viewed ~20 hours after treatment. (E, E’) Washington strain (PS1843) exposed to ZTDO viewed ~20 hours after treatment. (F, F’) Japan strain (RS5194) exposed to ZTDO viewed ~20 hours after treatment. Arrow indicates a possible embryonic plasma membrane. (G, G’) La Reunion strain (RSB001) exposed to ZTDO viewed immediately after treatment. (H, H’) Bolivia strain (RS5278) exposed to ZTDO viewed immediately after treatment. FM4-64 fluorescence images were taken at Intensity of 2 and exposure of 1 second. The scale bar of 20 μm in (A’) represents the same magnification for all images.

Next, we asked if the natural variation in ZTDO sensitivity is correlated with ZTDO-induced permeabilization of the eggshell. We found that like wild-type California PS312, late stage embryos of sensitive strains from Washington (PS1843) and Japan (RS5194) remain permeable to FM4-64 (Fig. 7E’-F’). Although the cytoplasmic staining in the Japan strain is weaker than PS312, we discovered that the embryos show a more severe abnormal cellular morphology than PS312. Almost no treated Japanese embryos ever reach the 2-fold or 3-fold stages; a majority arrest in gastrulation and the bean stage, perhaps as a result of more extensive tissue damage. In contrast, partially ZTDO resistant strains from La Reunion (RSB001) and Bolivia (RS5278) are largely impermeable to FM4-64, even when stained right after ZTDO exposure (Fig. 7G’-H’, compare to 7A’). Not all ZTDO susceptible embryos, however, show cytoplasmic staining by FM4-64. Whereas 100% of PS312, Washington, and Japan embryos do not hatch after a 4-hour ZTDO treatment (Fig. 3), only 65-83% of the embryos show permeable eggshells (Table 1). Similarly, whereas ~40% of Bolivia embryos do not hatch after ZTDO treatment, only 10% of the embryos show permeable eggshells. Only the *P. pacificus* La Reunion strain and *C. elegans* N2 show staining rates commensurate with ZTDO sensitivity at 10% and 0%, respectively. Also consistent with this trend, embryos of the ZTDO hypersensitive mutant, *Ppa-obi-1(tu404)* (Cinkornpumin et al., 2014), is more likely to show FM4-64 staining than wild-type PS312. Because the molecular weight of ZTDO is lower than FM4-64 (210 vs 608), ZTDO may enter openings that would size-restrict FM4-64. Yet it is not yet clear if ZTDO disrupts and penetrates the eggshell, or if disruption of the eggshell alone is sufficient to abolish embryogenesis. These results show that ZTDO can arrest development in part by increasing eggshell permeability and osmotic sensitivity, although ZTDO may also, in addition, damage embryonic tissue directly without detectable damage to the eggshell

**Table 1.** Staining of ZTDO-exposed 4-hour old embryos with the lipophilic membrane dye FM4-64. ^Viewed at 24 hours.

|  | **Strain** | **No. Trials**<br>**(sample**<br>**size)** | **Time**<br>**Viewed**<br>***ex utero*** | **Embryos**<br>**stained** | **Unhatched**<br>**embryos^** |
| --- | --- | --- | --- | --- | --- |
| Susceptible | PS312 (California) | 3(50) | 5 | 65% | --- |
|  | PS312 (California) | 3(50) | 24 | 83% | 100% |
|  | PS312 *obi-1(tu404)* | 3(50) | 24 | 90% | 100% |
|  | PS1843 (Washington) | 1(25) | 24 | 72% | 100% |
|  | RS5186 (Japan) | 2(35) | 24 | 77% | 100% |
| Resistant | RS5278 (Bolivia) | 2(50) | 5 | 10% | 40% |
|  | RS5419 (La Reunion) | 2(50) | 5 | 10% | 60% |
|  | *C. elegans* N2 | 2(50) | 5 | 0% | --- |
|  | *C. elegans* N2 | 2(50) | 24 | 0% | 0% |

## Discussion

Unlike broad-spectrum nematocides such as ivermectin, aldicarb, and monepantal that are administered in liquid or on solid media (Donnelly et al., 2013; Fru and Puoti, 2014; Goldman et al., 2007; Kampfe and Schutze, 1995; Wever et al., 2015), ZTDO is a synthetic version of the host Oriental beetle pheromone that exerts lethality as a volatile compound on *P. pacificus* embryos. Sensitivity to ZTDO appears to be the highest between the gastrula and tadpole stages, when two hours of odor exposure can prevent ~80% of embryos from hatching. A 3-hour exposure during the 4-8 hour period after synchronization is sufficient to achieve lethality, whereas a 4-hour exposure between the 0-4 hour period is necessary to arrest all embryos. ZTDO-exposed embryos arrest in various stages of incomplete embryogenesis that extend pre-hatching development from the usual 24 hours to 48 hours at 20°C, while at the same time preventing the aberrant embryos from hatching indefinitely. Lethality caused by ZTDO exposure results in a spectrum of incomplete embryogenesis, with the most severe phenotype found in the Japan embryos that never progress to the active zombryo stage, to the less severe zombryo phenotype of elongated, active embryos with partial formation of pharyngeal-like or gut-like lumens in the wildtype PS312 embryos. Because the most susceptible *P. pacificus* strain comes from the oriental beetle in Japan, in contrast to the La Reunion and Bolivia strains from other beetle species, ZTDO susceptibility in natural populations may have a role in maintaining host-parasite interactions.

Natural variation in ZTDO sensitivity is also correlated with ZTDO-induced permeabilization of the eggshell, although the percentage of embryonic staining is lower than the percentage of embryonic lethality. Since ascarosides are highly diverse in different species of nematodes and ascaroside-derived ascaroside glycolipids are required for eggshell assembly (Olson et al., 2012), it is possible that the lipid-like ascaroside side chain types and stoichiometry in the eggshell interact with the lipid-moiety of ZTDO to contribute to the variations in ZTDO-induced permeability among different *P. pacificus* strains. To fully account for

ZTDO’s embryonic lethality however, zygotic factors may also be involved. Early embryonic development up to the 50-cell stage in *C. elegans* and *P. pacificus* appears to be nearly identical, although the cell cycle is longer in *P. pacificus* (Schulze and Schierenberg, 2011; Vangestel et al., 2008). Transcripts of MED- 1/2 GATA factors appear between the 4-6 cell stage in early *P. pacificus* embryos (pers. Comm. M.Maduro)(Maduro et al., 2007), indicating that the maternal-zygotic transition occur only one division later in *P. pacificus* compared to *C. elegans.* 2-8 cell embryos are exposed to ZTDO after an hour of egg-laying, when the oldest embryos are already 1-hour into development at the start of exposure. Thus, during the standard ZTDO exposure between 0-5 hours *ex utero,* it is likely that zygotic gene products expressed at this time also have a role in mediating ZTDO sensitivity. In wild-type *P. pacificus* PS312, post-embryonic susceptibility to volatile ZTDO results in paralyzed J2 and dauer larvae (Cinkornpumin et al., 2014). However, it is not known if the same factor that mediates ZTDO-induced larval paralysis also mediates embryonic lethality.

Comparisons between the embryonic lethality caused by ZTDO and another embryocide reveal key differences in mode-of-action. In *C. elegans,* C22 is a potent and species-specific embryocide that disrupts the oocyte-to-embryo transition in the maternal environment (Weicksel et al., 2016). C22 acts only through the parental hermaphrodites and targets oocytes just prior to fertilization, but has no effect on any other developmental stages, such as embryos, larvae, and adults. In contrast, ZTDO does not interfere with *P. pacificus* egg viability *in utero,* but ZTDO acts strongly on early embryos *ex utero* before the tadpole stage, as well as subsequent J2 and dauer larvae stages after hatching. C22- induced embryonic lethality requires upregulation of the transcription factor LET-607 involved in protein trafficking, which in turn likely regulates the proper secretion of proteins needed for correct *C. elegans* eggshell assembly. Thus while C22 interferes with the permeability barrier during eggshell formation, ZTDO appears to permeabilize eggshells that are already formed. Both ZTDO and C22 increase eggshell permeability, which likely contributes to embryonic lethality due to increased osmotic sensitivity. However, in the case of ZTDO, because the fraction of ZTDO-permeablized eggshell detected by FM4-64 staining is lower than the fraction of embryos susceptible to the lethal effects of ZTDO, hatching ability may be affected even when damage to the eggshell or embryonic tissue is not detectable by FM4-64 staining. Furthermore, C22-treated embryos seldom proceed beyond the 100-cell stage, whereas around half of the ZTDO-treated embryos reach the superficially elongated 2-fold and 3-fold stages and continue developing 24 hours pass the normal hatching time, albeit with severe developmental defects.

Comparisons between the embryonic lethality caused by ZTDO and another embryocide reveal key differences in mode-of-action. In *C. elegans,* C22 is a potent and species-specific embryocide that disrupts the oocyte-to-embryo transition in the maternal environment (Weicksel et al., 2016). C22 acts only through the parental hermaphrodites and targets oocytes just prior to fertilization, but has no effect on any other developmental stages, such as embryos, larvae, and adults. In contrast, ZTDO does not interfere with *P. pacificus* egg viability *in utero,* but ZTDO acts strongly on early embryos *ex utero* before the tadpole stage, as well as subsequent J2 and dauer larvae stages after hatching. C22- induced embryonic lethality requires upregulation of the transcription factor LET-607 involved in protein trafficking, which in turn likely regulates the proper secretion of proteins needed for correct *C. elegans* eggshell assembly. Thus while C22 interferes with the permeability barrier during eggshell formation, ZTDO appears to permeabilize eggshells that are already formed. Both ZTDO and C22 increase eggshell permeability, which likely contributes to embryonic lethality due to increased osmotic sensitivity. However, in the case of ZTDO, because the fraction of ZTDO-permeablized eggshell detected by FM4-64 staining is lower than the fraction of embryos susceptible to the lethal effects of ZTDO, hatching ability may be affected even when damage to the eggshell or embryonic tissue is not detectable by FM4-64 staining. Furthermore, C22-treated embryos seldom proceed beyond the 100-cell stage, whereas around half of the ZTDO-treated embryos reach the superficially elongated 2-fold and 3-fold stages and continue developing 24 hours pass the normal hatching time, albeit with severe developmental defects.

ZTDO is a naturally occurring beetle compound that could deter reproductive growth of its associated *P. pacificus* while the beetle host is alive, and may be a lever for attenuated antagonism to keep necromenic nematodes from becoming pathogenic in a evolutionary detente (Sudhaus, 2009; Thompson, 2005). Because the physiological concentration of ZTDO that wild isolates of *P. pacificus* is likely to encounter is unknown, it is also possible that lower concentrations of ZTDO help to prolong rather than subvert embryonic development, as a way to coordinate with the death of the host beetle when bacterial food becomes imminent. In this respect, ZTDO-induced embryonic arrest may be analogous to host mediated hatching behavior found in the plant parasitic nematode *Heterodera glycines,* whereby unidentified root exudate from fresh samples stimulate hatching (Perry, 2002; Tefft and Bone, 1985; Thapa et al., 2017). However, whereas host cues increased hatching rate but has no effect on pre-hatching development in *H. glycines* (Thapa et al., 2017), the early exposure and terminal embryogenesis suggest ZTDO affects hatching rate indirectly by disrupting pre-hatching development, rather than the mechanisms required hatching. Since rapid evolution of host pheromone-induced embryonic lethality can be an important mechanism for facilitating host switching, we speculate that the natural variation in susceptibility to ZTDO is due to factors that is fast evolving at the population level. In the future, identification of the direct targets of ZTDO in the eggshell and embryonic tissue will lead to important insight into how the strain-specific embryonic response is achieved. Genetic screens for suppressors of the embryonic lethality will likely identify key factors mediating ZTDO recognition and pathway activation.

## Abbreviations

ZTDO: (Z)-7-tetradecen-2-one
ETDA: (*E*)-11-tetradecenyl acetate

## Acknowledgement

We thank B. Mojica and D. Kazerskiy for technical assistance.

## Author Contributions

TR and RLH designed the research, performed the experiments, analyzed data and wrote the manuscript.

**SI Fig. 1A. A two-fold ‘zombryo’ at 26 hours.** A two-fold ‘zombryo’ at 26 hours *ex utero* displays sporadic muscle contractions, but has not developed visible gut or pharyngeal-like structures.

**SI Fig. 1B. A two-fold ‘zombryo’ at 48 hours.** A two-fold ‘zombryo’ remains active 48 hours *ex utero* but is unable to hatch. The ZTDO-treated embryo has developed a midbody lumen, in contrast to the two-fold ‘zombryo’ at 26 hours.

**SI Fig. 1C. Two peanut stage ‘zombryos’ at 48 hours.** Two peanut ‘zombryos’ have developed gut lumens and muscle tissue by 48 hours *ex utero.* The embryos are unable to hatch but are active 24 hours passed the normal hatching time.

